# Biofilm/Persister/Stationary Phase Bacteria Cause More Severe Disease Than Log Phase Bacteria – I Biofilm *Borrelia burgdorferi* Not Only Display More Tolerance to Lyme Antibiotics But Also Cause More Severe Pathology In a Mouse Arthritis Model: Implications for Understanding Persistence, PTLDS and Treatment Failure

**DOI:** 10.1101/440461

**Authors:** Jie Feng, Tingting Li, Yuting Yuan, Rebecca Yee, Ying Zhang

**Affiliations:** Department of Molecular Microbiology and Immunology, Bloomberg School of Public Health, Johns Hopkins University, Baltimore, MD21205, USA

**Keywords:** *Borrelia burgdorferi*, stationary phase, variant forms, persisters, biofilm, arthritis

## Abstract

Lyme disease, caused by *Borrelia burgdorferi,* is the most common tick-borne illness in US and Europe. While most patients can be cured with a 2-4 week antibiotic therapy, about 10%-20% patients continue to suffer persistent symptoms of fatigue, pain or joint and muscle aches, and neurocognitive despite the treatment, a condition called post-treatment Lyme disease syndrome (PTLDS). The cause for PTLDS is unclear but one possibility is persistent infection with *B. burgdorferi. B. burgdorferi* is known to develop morphological variant forms such as round bodies and aggregated biofilm-like microcolonies as a log phase culture consisting of spirochete form grows into stationary phase. Here we isolated biofilm-like microcolony and planktonic form (spirochetal forms and round body) from stationary phase culture and found that the stationary phase planktonic form (SP) and microcolony form (MC) were much more tolerant to the current antibiotics for Lyme disease, doxycycline, ceftriaxone and cefuroxime than log phase spirochete form (LOG). In addition, we also compared the ability of the variant forms to cause disease in a mouse arthritis model. Surprisingly, the MC in particular and the SP caused a more severe arthritis with an earlier onset of inflammation and joint swelling than LOG. MC-infected mice showed significant joint swelling as early as 9 days post-infection, while the LOG and SP did not cause significant swelling. At 21 days, the joint swelling of the MC group dramatically increased and peaked, while the SP showed significant swelling at this time but less severe than the MC group. The LOG infected mice were just beginning to develop joint swelling at 21-day post-infection, with only slight swelling. At 30-day post infection, the SP group mice also developed similar severity of joint swelling as the MC group, but the LOG group still did not show significant swelling. However, at 35-day post infection, all three infected groups showed similar degree of significant joint swelling. Thereafter, the joint swelling of the three infected groups waxed and waned during the 90-day observation. Thus, we established a new biofilm-inocula mediated visual arthritis model that could facilitate more efficient evaluation of treatment regimens for persistent *B. burgdorferi* infections. Our findings provide new insight about disease pathogenesis and may have implications for understanding PTLDS and PTLDS treatment failure, due to possible biofilm inoculation during tick-bite. This biofilm/persister seeding model may be valid for different microbial infections and facilitate developing more effective treatments of persistent infections in general.

## Introduction

Lyme disease (LD) is the most prevalent tick-borne illness and an important emerging zoonosis in the United States with an estimated 300,000 cases per year [1]. The causative agents of Lyme disease are pathogenic Borrelia species including *B. burgdorferi, B. afzelii, B. garinii*. *B. burgdorferi* is the predominant cause of human LD in North America. In humans, Lyme disease may cause a local erythema migrans (EM) rash at the site of the tick bite and then readily disseminates through the bloodstream to other tissues, setting up an infection that can last for months to years. Patients with Lyme disease are routinely treated with doxycycline, amoxicillin or cefuroxime, which effectively hastens the resolution in most cases. However, about 10%-20% patients continue to suffer lingering symptoms of fatigue, pain or joint and muscle aches, and neurocognitive manifestations that last 6 months or more despite treatment, a condition called “post-treatment Lyme disease syndrome” (PTLDS) [2]. While the cause of PTLDS is complex and remains to be determined, one of the possible explanations is persistent *B. burgdorferi* infection due to persisters not killed by the current antibiotic treatment, which evade host immune clearance and drive immunological responses continually as shown in various animal models [3-6]. A recent study in humans demonstrated the recovery of *B. burgdorferi* DNA by xenodiagnoses in patients despite antibiotic treatment [7]. Findings indicate that current Lyme disease treatment may not sufficiently eliminate *B. burgdorferi* persisters or that the immune system fails to clear persisting organisms or bacterial debris, which may be the underlying cause for those who suffer from unresolved Lyme disease symptoms. In contrast to other bacterial pathogens that cause persistent infections such as *M. tuberculosis* and *E. coli,* an unusual feature of the in vivo persistence of *B. burgdorferi* is the lack of culturability of the persisting organisms despite the demonstration of its DNA and even increased DNA content by PCR or by xenodiagnosis [4, 5]

In addition to the above in vivo persistence, *B. burgdorferi* has recently been shown to develop persisters in vitro in cultures as shown by tolerance to current Lyme antibiotics doxycycline, amoxicillin and cefuroxime [8-10]. In addition, *B. burgdorferi* can develop morphological variants including spirochetal form, round body form, or cystic form, and aggregated biofilm-like microcolonies as the culture grows from log phase to stationary phase or under stress conditions in vitro [8, 11]. The variant forms such as cystic and round body forms have been found in vivo in brain tissues of Lyme borreliosis patients [12] but their role in persistent form of the disease is controversial [13]. We showed that stationary phase cultures contain different morphological variants including planktonic spirochetal form, round body form and aggregated microcolony form [8, 11], which have varying levels of persistence or can be considered different types of persisters in comparison to the log phase culture which mainly consists of growing spirochetal form with no or few persisters [8]. However, the previous study showing different degree of tolerance to antibiotics by the variant forms was performed with mixed stationary phase culture [8], but only individual isolated variant forms. In addition, although different variant forms of *B. burgdorferi* seem to elicit different host cytokine response [14], their ability to cause disease has not been evaluated. In this study, we isolated three different forms of *B. burgdorferi*, including growing log phase spirochetes (LOG), non-growing stationary phase planktonic (SP) form, and aggregated biofilm-like microcolony (MC) form and compared them in treatment with different antibiotics in vitro and found indeed the MC form and SP form to be more tolerant to antibiotics. More importantly, we found that the MC form and the SP form caused a more severe arthritis in mice than the log phase spirochetal form.

## Results

### Separation of aggregated microcolony form from planktonic form in stationary phase culture

Previous studies demonstrated that *B. burgdorferi* develops multiple morphological forms including spirochetal form, round bodies (cysts), and aggregated microcolony form [11, 12]. Log phase *B. burgdorferi* culture is mainly in spirochetal form, but stationary phase culture is dominated by coccoid or round-body forms and aggregated micro-colony forms [11]. We showed that the current clinically used antibiotics used for treating Lyme disease had high activity against log phase *B. burgdorferi* but had very limited activity against the stationary phase cells [11]. We also demonstrated that aggregated microcolony forms are more tolerant to antibiotics than planktonic spirochetal and round body forms in the stationary phase culture [8]. To identify proteins that are preferentially expressed in the variant forms to shed new light on persistence mechanisms, here we first separated the aggregated microcolony form from the planktonic form (spirochetal and round body) in stationary phase *B. burgdorferi* culture according to their different densities using low speed centrifugation. After multiple differential centrifugation separation, microcolony form and planktonic form were separated effectively (Figure 1B and C). Consistent with our previous study [8], here we found that the 5 day old log phase cells were mostly in spirochetal form (Figure 1A), while the 10-day old stationary phase culture contained, in addition to aggregated biofilm-like microcolonies (Fig. 1B), planktonic forms made up of not only spirochetes but also many round body cells (Figure 1C).

**Figure 1.**
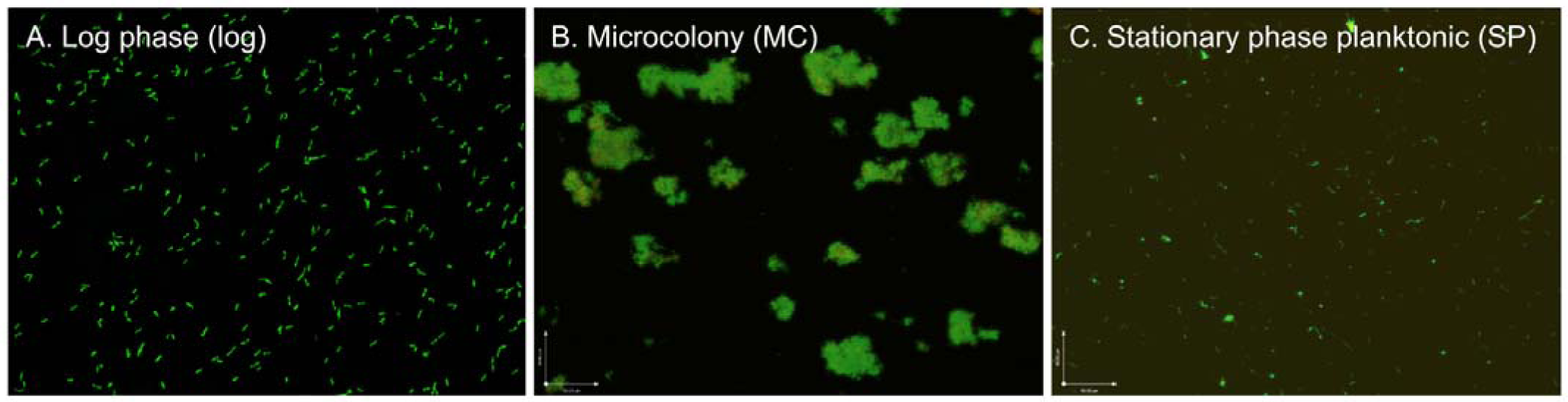
Representative images of a 3-day old log phase (A), 10-day old stationary phase microcolony form (B) and 10-day stationary phase planktonic form (C) of *B. burgdorferi.* Cells were stained with SYBR Green I/PI and observed using fluorescence microscopy at 40 X magnification.

### Different B. burgdorferi forms have different drug susceptibility in vitro

Since our previous studies showed that stationary-phase *B. burgdorferi* cells are more tolerant to antibiotics [8, 11], here we compared the drug susceptibility between the stationary phase planktonic form, aggregated microcolony form and the log phase *B. burgdorferi*. After 7-day drug treatment, the stationary phase planktonic (SP) form and microcolony form (MC) *B. burgdorferi* were found to be much more tolerant to the current clinically used antibiotics, doxycycline, ceftriaxone and cefuroxime (5 μg/ml) (Figure 2). The residual viability after drug treatment showed statistically significant difference between the log phase form and two stationary phase forms SP and MC (Figure 2A). By microscope checking, we found that most of the log phase spirochetal *B. burgdorferi* cells were eradicated by the 7-day doxycycline or ceftriaxone treatment. In contrast, most of the stationary phase planktonic and microcolony form of *B. burgdorferi* cells were still alive as shown by green fluorescence after the 7-day doxycycline or ceftriaxone treatment (Figure 2B).

**Figure 2.**
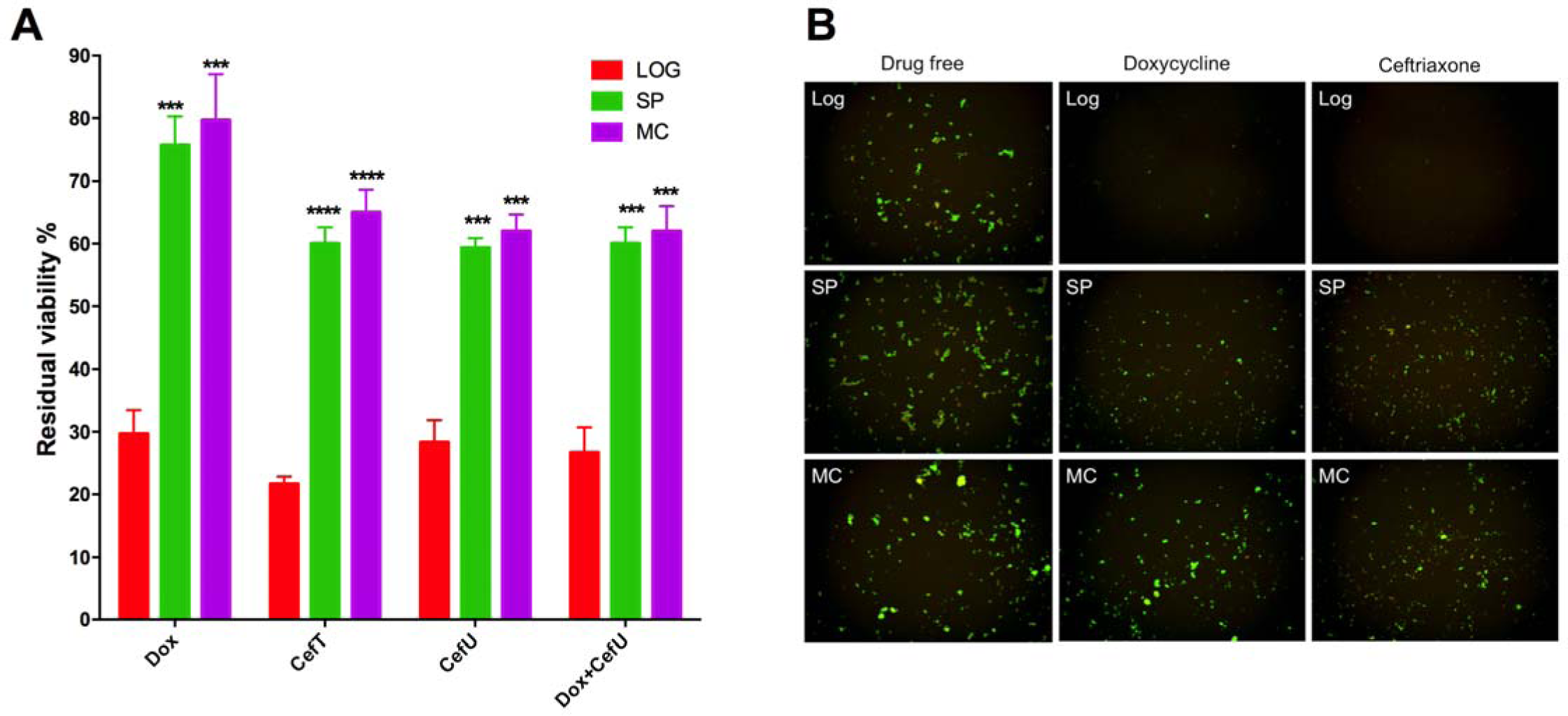
Susceptibility of log phase (LOG), stationary phase planktonic (SP) and microcolony form (MC) *B. burgdorferi* to 5 µg/mL clinically used antibiotics after 7-day treatment. (A) The percentages of residual live cells (means ± standard deviation) were determined with SYBR Green I/PI assay. The asterisks indicate significant susceptibility difference (triple asterisks, *P <* 0.001; quadruple asterisks, *P <* 0.0001) as compared to the log phase group. (B) Representative images of the different form *B. burgdorferi* treated with doxycycline and ceftriaxone for 7 days followed by staining with SYBR Green I/PI assay (40 X magnification).

### Stationary phase planktonic form and microcolony form of B. burgdorferi cause more severe persistent Lyme arthritis than the log phase spirochete form

To investigate the effects of different forms of *B. burgdorferi* on development of arthritis in the mice, C3H/HeN mice were infected by subdermal injection with 10^5^ cells of N40 strain. All injected mice became infected, as determined by ear punch and bladder culture at the end of experiments. During the 90-day infection period, we used caliper measurements to monitor the swelling or edema of ankle and knee joint, which can directly reflect the degree of the inflammatory response to *B. burgdorferi* infection [15]. We found that the different forms of *B. burgdorferi* showed different effects on inducing ankle joint swelling in C3H/HeN mice, especially in the early stage of 21 days (Figure 3). The microcolony form of *B. burgdorferi* (MC) showed most obvious pathogenic effect in terms of the ability to cause joint swelling compared to the other forms and uninfected control (Figure 3). MC infected group showed significant swelling (*p<*0.005) compared with uninfected group as early as 9 days (Figure 3A), however, the log phase spirochete form and stationary phase planktonic form did not cause significant swelling (*p>*0.05) (Figure 3A). At 21 days, the joint swelling of the MC group dramatically increased and reached peaked (Figure 3); but the SP form showed significant swelling (*p<*0.005) at this time point but less severe than the MC form infected group (Figure 3). The log phase *B. burgdorferi* infected mice were just beginning to develop joint swell at Day 21-post infection, with only slight swelling (Figure 3A and C). At 30-day post infection, the SP group mice were also found to develop similar severity of joint swelling as the MC group, but the log phase group still did not show significant swelling (*p>*0.05) (Figure 3A). However, at 35-day post infection, all three infected groups showed similar degree of significant joint swelling (*p<*0.05) compared to the uninfected control group (Figure 3A and C). After 35-day infection, the joint swelling of the three infected groups relapse during the 90-day observation, while the severity of ankle joint swelling did not have significant difference between the three different *B. burgdorferi* infected groups post 35-day infection (Figure 3B).

**Figure 3.**
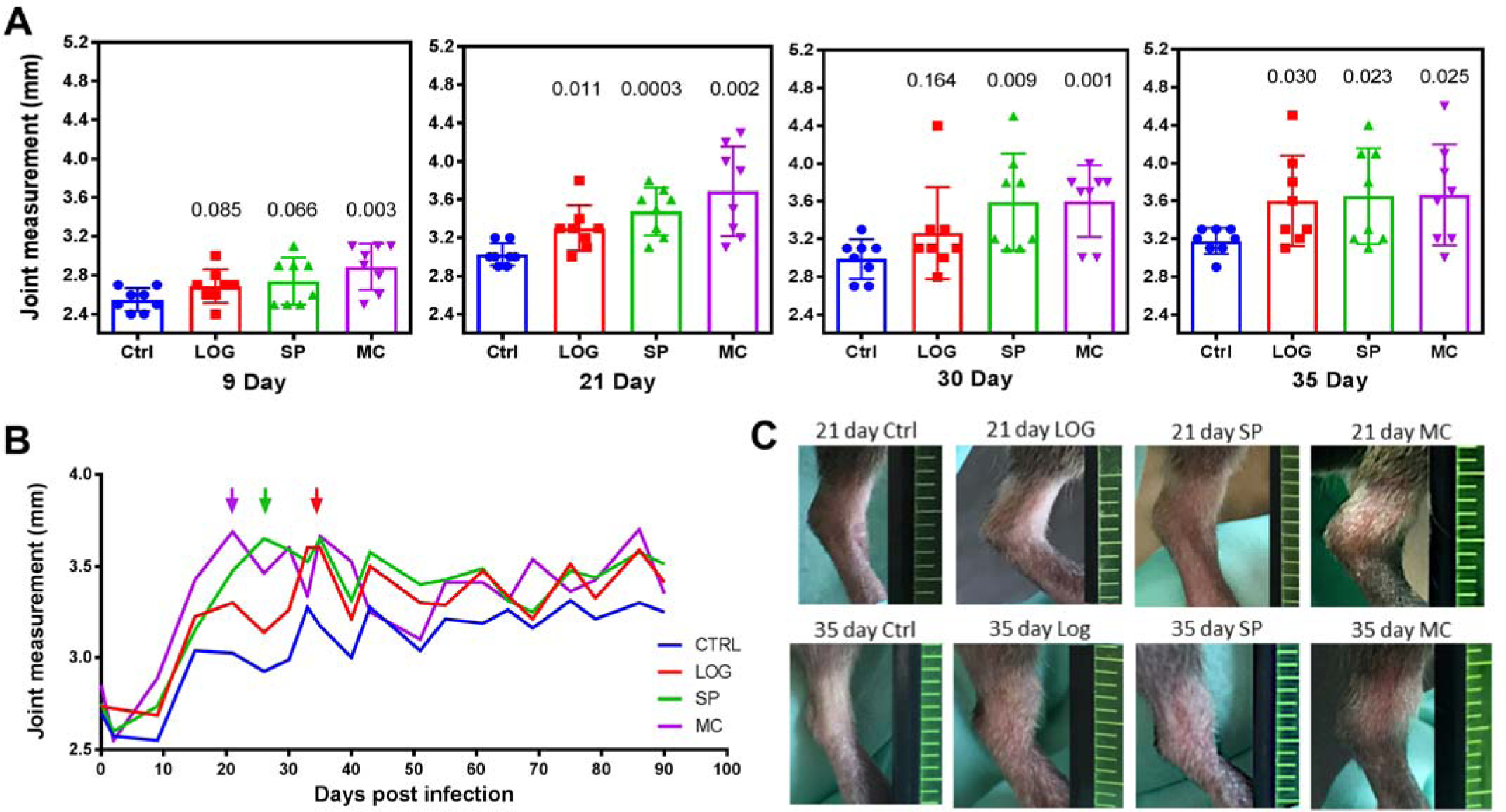
Ankle joints measurements of different form *B. burgdorferi* infected and uninfected C3H/HeJ mice. Measurements of every ankle joint were made for each mouse every week for 90 days. (A) At early stage infection (before 35 day), the group of microcolony form (MC), stationary phase planktonic form (SP) and log phase form (LOG) infected mice showed different severity on ankle swelling. Measurement values, means and standard deviations (error bars) were shown for each group. Statistical analyses were performed using the standard *t-*test, comparing the joint measurements of infected and uninfected groups. (B) ankle joint swelling mean values of different groups during the 90-day measurements. The colored arrows indicate the earliest time of reaching the most severely swelling level of the corresponding groups. (C) The images of the most severe swollen ankles in different groups at 21 day and 35 day, respectively.

## Discussion

In this study, we show that different variant forms of *B. burgdorferi* are different types of persisters with varying degree of persistence and are not killed readily by current Lyme antibiotics (Fig. 2). However, the significance of the in vitro persisters to in vivo was not clear. Here, we found that the variant forms MC and stationary phase planktonic form (SP) caused more severe disease (arthritis) than the log phase spirochetal form of *B. burgdorferi* (Fig. 3). In addition, it is known that *B. burgdorferi* in the tick could develop variant forms that could represent different persisters [16]and that biofilm bacteria have been found in skin biopsies in humans [17]. Previously, we hypothesized that persistent bacterial infections could be caused by “persister-seeding” theory [18]. In view of these observations and in particular our new findings in this study, we propose a new theory of PTLDS to better explain the persisting symptoms despite the standard Lyme antibiotic treatment: biofilm “seeding” theory. This theory can be called “Inocula-Dependent Severity of Disease” determined by the status/quality of the inocula bacteria, ranging from spirochetes to round bodies, microcolonies and biofilms. Quality of the inocula persisters matter, that is, the outcome of the infection is to a large extent dependent on the quality of the inoculum. A small inoculum may be sufficient to cause disease if it is mainly microcolony or persister/mother cell or stem-like cell. Persistent infections can be caused by *B. burgdorferi* morphologic variant forms, different types of persisters (spirochetal form, round body, cyst, microcolony, biofilm), and that different forms may be able to cause different disease severity, with spirochetal form causing mainly active or acute form of the disease that is more easily curable, while round bodies and microcolonies, and biofilms may cause more severe disease, given the healthy competent host immune response. This new model of biofilm and persister bacteria seeding explains some mysterious features of PTLDS, i.e., inability of the current Lyme antibiotics to eradicate or cure the 10-20% patients who fail the initial standard 2-4 week antibiotic treatment; early onset of the PTLDS or continued PTLDS despite early and timely treatment. Thus, infection caused by biofilm/persister bacteria will not respond well to the current standard Lyme antibiotics, which is essentially a treatment failure. On the other hand, there is another type of persistent disease or PTLDS, i.e., the initial infection is not diagnosed and treated in time such that more persisters and biofilm develop leading to a persistent infection that is more difficult to treat or cure. Thus, although PTLDS can be a very heterogeneous condition, we can broadly divide PTLDS into two types: Type I PTLDS, or early development persistent disease where persister/biofilm inocula-dependent with severe disease from the beginning; our biofilm microcolony induced mouse arthritis model (Zhang-Feng model) is a representative of this Type I PTLDS; and Type II PTLDS, or late development persistent disease, where the long term mouse persistence model which allows an acute infection to develop into a persistent phase of the disease through prolonged infection before treatment (Barthold-Hodzic model). The development of this biofilm-inocula seeding model or Type I PTLDS model may allow us to more efficiently evaluate persister drug regimens which we found to be effective in vitro [8] and have implications for better understanding disease pathogenesis and developing better diagnostics and more importantly better treatments for PTLDS.

The observation that biofilm/persister inocula produce a more severe disease than log phase growing bacteria as shown here has not been paid attention to previously in the field of bacterial pathogenesis studies. It is generally assumed that all bacteria are created equal and are equally able to cause disease in a host. Most studies use log phase bacteria for infections and pay more attention to the number of bacteria used for infection rather than the quality of the bacterial inocula[19-22]. Our findings that biofilm/persister inocula produce a more severe disease than log phase bacteria call attention to quality of the bacteria inocula used for infection being an important consideration in disease model and a re-evaluation of the possible differences in pathogenesis and host response associated with the more severe form of the disease induced by the biofilm/persister inocula.

The mechanisms by which the biofilm/persister inocula produce a more severe disease remain to be determined. Several possibilities exist. The genes and pathways turned on in biofilm/persister inocula [18] may allow the persister bacteria to survive the attack by the host immune system such that they can better establish the infection and cause disease. In contrast, log phase bacteria may be killed more easily by the immune systems such that they do not cause as severe disease as the biofilm/persister inocula. Indeed, it has been shown that *B. burgdorferi* persisters tolerant to antibiotics express more virulence factors such as decorin-binding proteins (DbpAB), CRASP BB_A68, ErpQ, BdrEFMRVW, BB_I29, Clp protease, and DNA repair proteins in RNA-seq analysis [23] and also in proteomic studies (Feng et al., to be published). Thus, there is a molecular basis for the increased virulence of the biofilm/persister inocula in the case of *B. burgdorferi*.

The biofilm/persister seeding model [18], where the biofilm/persister bacteria cause more severe and persistent disease, has been shown to be valid not only for Borrelia infection in this study, but also for other bacterial infections such as *S. aureus* persistent infections (Yee R et al., see companion article). We believe this biofilm/persister seeding model to be valid for different microbial infections and will have broad implications for understanding disease pathogenesis and for developing more effective treatments of persistent infections and even cancer in general.

## Materials and Methods

### Strain, media, and culture techniques

The low passaged (less than 5 passages) *Borrelia burgdorferi* strain B31 5A19 and strain N40 were kindly provided by Dr. Monica Embers [9] and Dr. Emir Hodzic. *B. burgdorferi* strains were grown in BSK-H medium (HiMedia Laboratories Pvt. Ltd.) with 6% rabbit serum (Sigma-Aldrich, St. Louis, MO, USA). All culture medium was filter-sterilized by 0.2 μm filter. Cultures were incubated in microaerophilic incubator (33°C, 5% CO_2_) without antibiotics.

### Antibiotics and drug susceptibility assay

Antibiotics including doxycycline, ceftriaxone and cefuroxime were purchased from Sigma-Aldrich (St. Louis, MO, USA) and dissolved in water at appropriate concentrations to form stock solutions. All antibiotic stocks were filter-sterilized using a 0.2 μm filter. The residual viability of *B. burgdorferi* cells treated with antibiotics or drug combinations were evaluated using the SYBR Green I/PI assay combined with fluorescence microscopy as described previously [11]. Briefly, the ratio of live and dead cells was calculated by the ratio of green/red fluorescence and the regression equation and regression curve with least-square fitting analysis.

### Separation and preparation of microcolony form and planktonic form from stationary phase culture

After incubation for 10 days, 1 ml stationary phase *B. burgdorferi* culture (∼10^7^ spirochetes/mL) was centrifuged at 800 × g for 10 min, and the supernatant was transferred to a new tube as stationary phase spirochetal form. The bottom 50 μl microcolony rich culture was resuspended and centrifuged at 800 × g 3 times to remove the planktonic spirochetes. The stationary phase spirochetal form and microcolony form were checked with fluorescence microscope to ensure their morphologies before being used for the infection and mass spectrum analysis (see below).

### Animals and infections

5-6-week-old female C3H/HeN mice purchased from Charles River were infected by subdermal inoculation of the dorsal thoracic midline with about 10^5^ cells of different forms of *B. burgdorferi* strain N40, i.e. log phase spirochetal form, stationary phase planktonic form, and stationary phase aggregated biofilm-like microcolony form. Infection was confirmed by the culture of *B. burgdorferi* from the ear punch and bladder. The infection was followed over a period of 90 days to monitor the degree of arthritis induced by different morphological forms. All experiments were performed in accordance with the Animal Care and Use Committee of Johns Hopkins University.

### Assessment of arthritis severity

Caliper measurements of ankle and knee joints were performed every two days during the infection by an investigator blinded to the experimental groups. For assessment of histopathologic lesions, the heart, the skin and tissue of rear ankle joints were separately fixed in 10% neutral buffered formalin (Sigma, USA). Fixed tissues were decalcified and embedded in paraffin, sectioned at 3 mm, and stained with H&E.

Lesions were scored in a blinded fashion, with each slide receiving a score of 0–5 for the characteristics of the disease such as polymorphonuclear leukocyte and mononuclear cell (lymphocytes, monocytes, macrophages) infiltration into inflammatory processes, tendon sheath thickening (hypertrophy and hyperplasia of surface cells and/or underlying dense sheets of cells resembling immature fibroblasts, synoviocytes, and/or granulation tissue), reactive/reparative responses (periosteal hyperplasia and new bone formation and remodeling), and overall lesion (composite score based on all lesions observed in six to eight sections per joint), with 5 representing the most severe lesion, and 0 representing no lesion [24].

## Acknowledgments

We thank Monica Embers and Emir Hodzic for providing borrelia strains, and Tim Sellati for helpful discussion. We acknowledge the support by Global Lyme Alliance, LivLyme Foundation, NatCapLyme, and the Einstein-Sim Family Charitable Fund.

